# Unraveling Honey Bee’s Waggle Dances in Natural Conditions with Video-Based Deep Learning

**DOI:** 10.1101/2024.11.16.623982

**Authors:** Sylvain Grison, Rajath Siddaganga, Shrihari Hegde, James Burridge, Pieter M. Blok, Smitha Krishnan, Axel Brockmann, Wei Guo

## Abstract

1. Wild and managed honey bees, crucial pollinators for both agriculture and natural ecosystems, face challenges due to industrial agriculture and urbanization. Understanding how bee colonies utilize the landscape for foraging is essential for managing human-bee conflicts and protecting these pollinators to sustain their vital pollination services. To understand how the bees utilize their surroundings, researchers often decode bee waggle dances, which honey bee workers use to communicate navigational information of desirable food and nesting sites to their nest mates. This process is carried out manually, which is time-consuming, prone to human error and requires specialized skills.
2. We address this problem by introducing a novel deep learning-based pipeline that automatically detects and measures waggle runs, the core movement of the waggle dance, under natural recording conditions for the first time. We combined the capabilities of the action detector YOWOv2 and the DeepSORT tracking method, with the Principal Component Analysis to extract dancing bee bounding boxes and the angles and durations within waggle runs.
3. The presented pipeline works fully automatically with videos taken from wild *Apis dorsata* colonies in its natural environment, and can be used for any honey bee species. Comparison of our pipeline with analyses made by human experts revealed that our procedure was able to detect 93% of waggle runs on the testing dataset, with a run duration Root Mean Squared Error (RMSE) of less than a second, and a run angle RMSE of 0.14 radians. We also assessed the generalizability of our pipeline to previously unseen recording conditions, successfully detecting 50% of waggle runs performed by *Apis mellifera* bees from a colony managed in Tokyo, Japan. In parallel, we discovered the most appropriate values of the model’s hyperparameters for this task.
4. Our study demonstrates that a deep learning-based pipeline can successfully and automatically analyze the waggle runs of *Apis dorsata* in natural conditions and generalize to other bee species. This approach enables precise measurement of direction and duration, enabling the study of bee foraging behavior on an unprecedented scale compared to traditional manual methods contributing to preserving biodiversity and ecosystem services.

## 1 Introduction

Wild and managed bees provide important pollination services that are crucial for our food and nutritional security. Additionally, they are crucial in supporting diversity and resilience of managed and natural ecosystems and thereby aid a multitude of other associated ecosystem services (Krishnan et al., 2012; Potts et al., 2016; Warrit et al., 2023). While human-managed colonies of the Western honey bee, *Apis mellifera*, are the workhorses of industrial agricultural food production in Europe, America, Australia and to a lesser extent in Asia, a group of wild native honey bee species (*Apis dorsata*, *Apis laboriosa*, *Apis cerana*, and *Apis florea*) are the dominant pollinators in natural and agricultural habitats of Asia (Papa et al., 2022; Stewart et al., 2018). Commercial beekeeping for pollination service is only recently gaining popularity in a few parts of Asia while largely farmers rely on wild honey bees. Due to the large colony size and their massive food demands, the wild *Apis dorsata* likely is the most important pollinator in the Asian tropics. In natural habitats, the colonies build their nest on large trees or cliffs close to water sources, whereas in urbanized areas, they prefer to nest on tall, human-made structures such as water towers and overhangs of high-raised buildings including apartment balconies (Stewart et al., 2018; Walter & Brockmann, 2022). The urban human population, increasing due to rapid urbanization, fear the aggressive defensive attacks of these honey bees and often demands the eradication of colonies from highly populated public and private areas (Alex et al., 2017; Ramachandra et al., 2013; Vanjare et al., 2014; Walter & Brockmann, 2022).

In honey bees, workers communicate the location of food sources and nesting sites to each other via a stereotypical movement pattern called waggle dance (Dyer, 2002; Frisch, 1993). The waggle dance is composed of several straight forward runs, during which the bee shakes her abdomen, and left or right return loops which will lead the bee back to the starting position from where she will initiate another waggle run. Thus, a dancing bee communicates navigational information about the direction and distance of rich food sources to unemployed nestmates and helps to optimize the colony’s foraging efforts (Nürnberger et al., 2019; Seeley, 2011):

- The direction of the movement with respect to gravity indicates the flight directions: an upward direction (against gravity) indicates a location towards the direction of the sun, and a downward direction indicates a location opposite to the sun’s position.
- The duration of the waggle phase correlates with the flight distance.

Understanding how the colonies select foraging sites in various habitats, how they are affected by human activities (land use land cover change, agricultural activities, etc.) would help us to better safeguard their nesting and foraging needs and, in the process, to manage the human-bee conflict and protect honey bees to sustain their important pollination services (Couvillon et al., 2014a; Rutschmann et al., 2023; Young et al., 2021).

In practice, the foraging locations of honey bee colonies can be monitored via recording and decoding the waggle dance (Baensch et al., 2020; Beekman et al., 2015; Couvillon et al., 2014b; Samuelson et al., 2022). Traditionally, this process involves watching a video on a screen to identify where and when waggle run movements occur, reporting the number of video frames for each waggle run, measuring the direction of the movement between the first and the last position for each waggle run, and averaging these values over multiple waggle runs of same individual. The annotator needs to inspect the video multiple times to avoid missing any waggle run, and loop back continuously to set the waggle run’s beginning and ending with precision. Analyzing videos for hours while maintaining consistent focus is a laborious and time-consuming task that requires specialized skills. Several efforts have been made to generate semi- or fully automated procedure to analyze waggle dance orientation and direction from video clips. For instance, Rau (2014) used a pixel detector based on detecting waggle-like frequencies. Data were captured in a uniquely designed environment featuring an observation hive and 100fps cameras. Besides organizing specific recording conditions, their algorithm requires manual configuration of its parameters, such as the discriminatory intensity range used to classify dance pixels. These limitations reduce the algorithm’s adaptability to varying conditions and require expert knowledge to tune it properly. Also, the method is not robust to the dances that occur close to each other since it does not use visual features and would often merge them into one individual. Wario et al. (2015) detected waggle-like movements with a similar method based on bee shaking movement frequency. They also introduced a new deep learning method to detect dances in laboratory conditions (Wario et al., 2017). The latter method is based on four modules in series, the first of which is based on a waggle-like frequency detector.The first module’s reliance on empirically determined thresholds for run detection is a significant limitation, as it may not be optimal and impacts the performance of subsequent modules that depend on its output. This method also requires a specific environment featuring a high-frequency (120fps) camera with an observation hive illuminated by LEDs, which limits the application range. Saghafi and Tsokos (2017) used a dataframe of bee head angles regarding the rest of the body, of dancing and non-dancing bees from a a previous study. Training such a detector would require significant annotation time. Using an observational hive under experimental set up, Reece et al. (2020) used a detection method based on frame difference on 25-fps videos to detect bees with abnormal rapid movements. This method achieved a high precision of 92.8% but still missed many runs as the recall was 78.2 %. Bozek et al. (2021) and Kongsilp et al. (2024) developed a whole hive tracking method and mentioned about a potential waggle dance detector built as a downstream application of it. Implementing such a method would require a heavy annotation step to train the model. As illustrated above, previous works exhibit significant drawbacks. These include the need for specialized recording conditions (e.g., observation hives, high-speed cameras, artificial lighting), reliance on manually defined thresholds, limiting adaptability, inability to handle closely dancing bees, high false negative rates, and the requirement for extensive manual annotation for training machine learning models. Additionally, many methods struggle with generalizing to varied hive conditions and require significant expert knowledge for optimal setup and operation.

Thereby, despite the critical importance of understanding honey bee foraging behavior for ecological and agricultural purposes, current methods do not provide a well-rounded solution to analyze runs. Our study addresses these limitations by focusing on the following research question: How can deep learning approaches be optimized for the automatic detection and analysis of bee waggle runs across diverse environmental conditions without requiring specialized recording setups or context-specific parameter tuning? By bridging the gap between high-throughput video analysis with deep learning and ecological and behavioral studies, we built a comprehensive pipeline to automatically detect waggle runs in natural conditions, and extract key mapping parameters like duration and angle with a high accuracy using standard Full HD cameras, allowing for large-scale ecological studies by taking advantage to its ability to generalize to new conditions. The main contributions of this study are summarized as follows:

- The first fully automatic deep learning approach to detect bee waggle runs and extract their key parameters in videos taken in natural conditions, with limited data annotation.
- An in-depth analysis of the effect of different hyperparameters on the performance.
- The public release of our dataset and software.

## 2 Materials and methods

### 2.1 Training and evaluation dataset preparation

The colony of Apis dorsata used in the study was located in a primarily residential district in the northern part of the city of Bangalore, India. The recordings were made at 50 fps using commercially available video cameras (Panasonic HC-X929 and Panasonic HC-V707). The videos were recorded in Full HD (1920×1080 pixels) under natural conditions and the camera was placed at a distance of 0.75 m from the colony, ensuring that the entire colony was fully covered and individual waggle dances were clearly visible (Fig. 1). The recorded videos (N=5) were split into 187 clips (448x448 resolution), each containing one to three waggle runs from one or several different bees. These videos were allocated as follows: 127 clips for the training dataset (68%), 29 clips for the validation dataset (15%), and 31 clips for the testing dataset (17%). Each clip was annotated using the Darwin V7 annotation tool (https://www.v7labs.com/) to include the following information:

- Bounding boxes around the body of dancing bees, during the shaking phase only (not during the return run).
- Key points where the abdomen and thorax separate on dancing bee bodies, at the first and the last positions within runs. Each run angle was then measured as the angle of the line linking these two points, relative to the vertical axis of the hive, which is also the vertical axis with respect to gravity (Fig. 2).

**Figure 1:**
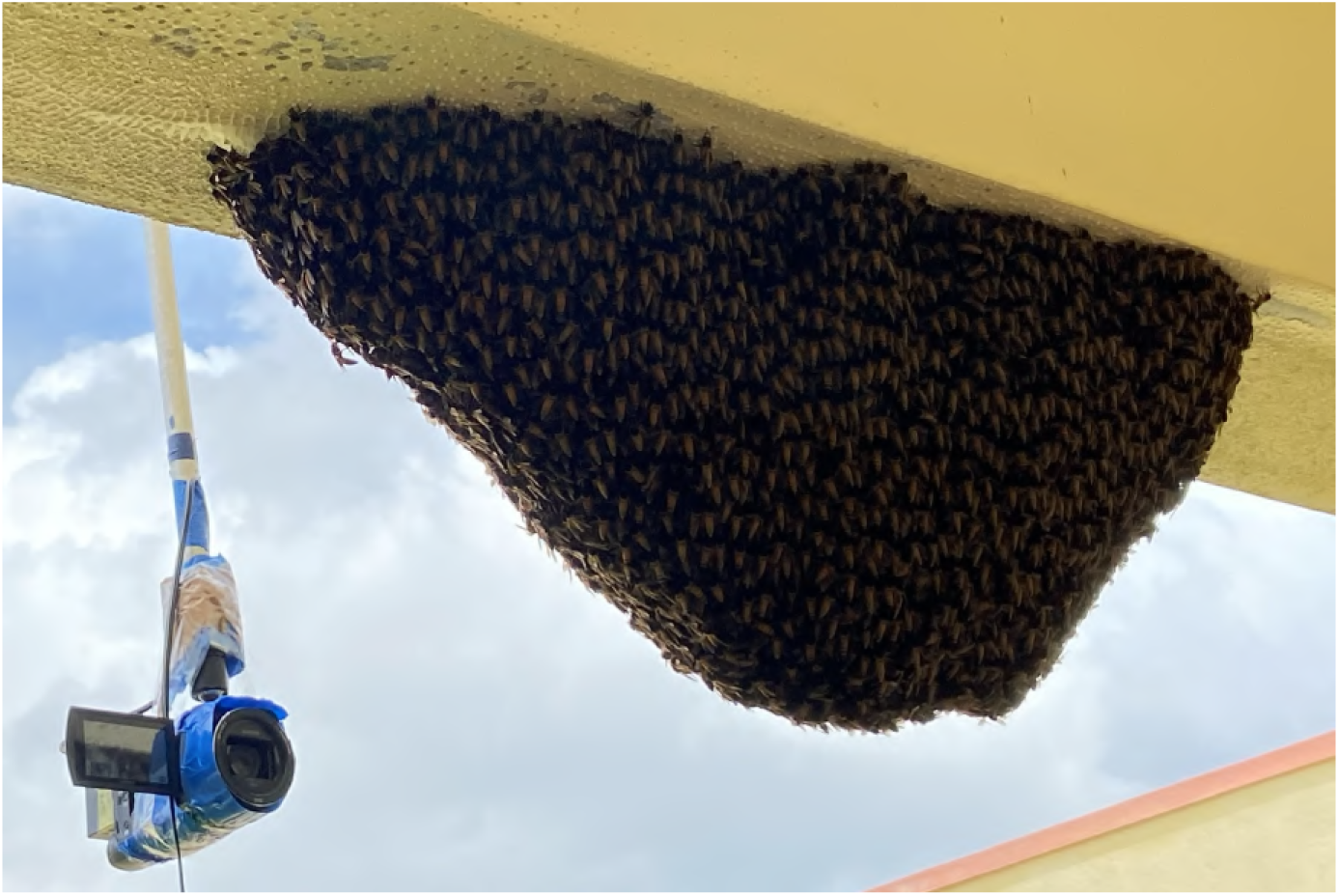
Photo of recording setup for capturing videos of waggle runs in a *Apis dorsata* hive under natural conditions.

**Figure 2:**
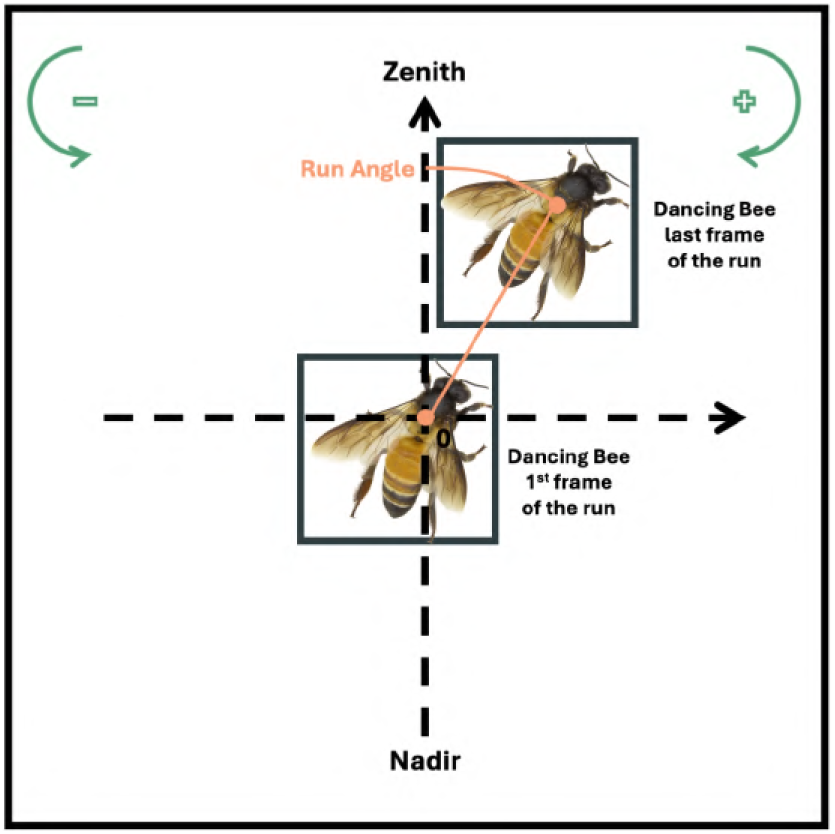
Angle measurement method in our experiments.

### 2.2 Waggle run detector

#### 2.2.1 Pipeline overview

In this study, we designed a pipeline to automatically analyse honey bee workers video clips taken in natural conditions by detecting waggle runs, extracting key parameters from them, and translating them to real-world coordinates. As shown in Fig. 3, our pipeline is made of two modules :

- The run recognition module takes a video as an input and outputs dancing bounding boxes proposals for each frame of the video.
- The run tracking module takes the bounding boxes output from the previous module and connects them over time to produce tubes, the equivalent of bounding boxes for a video. For each run, the direction and the duration are also extracted.

**Figure 3:**
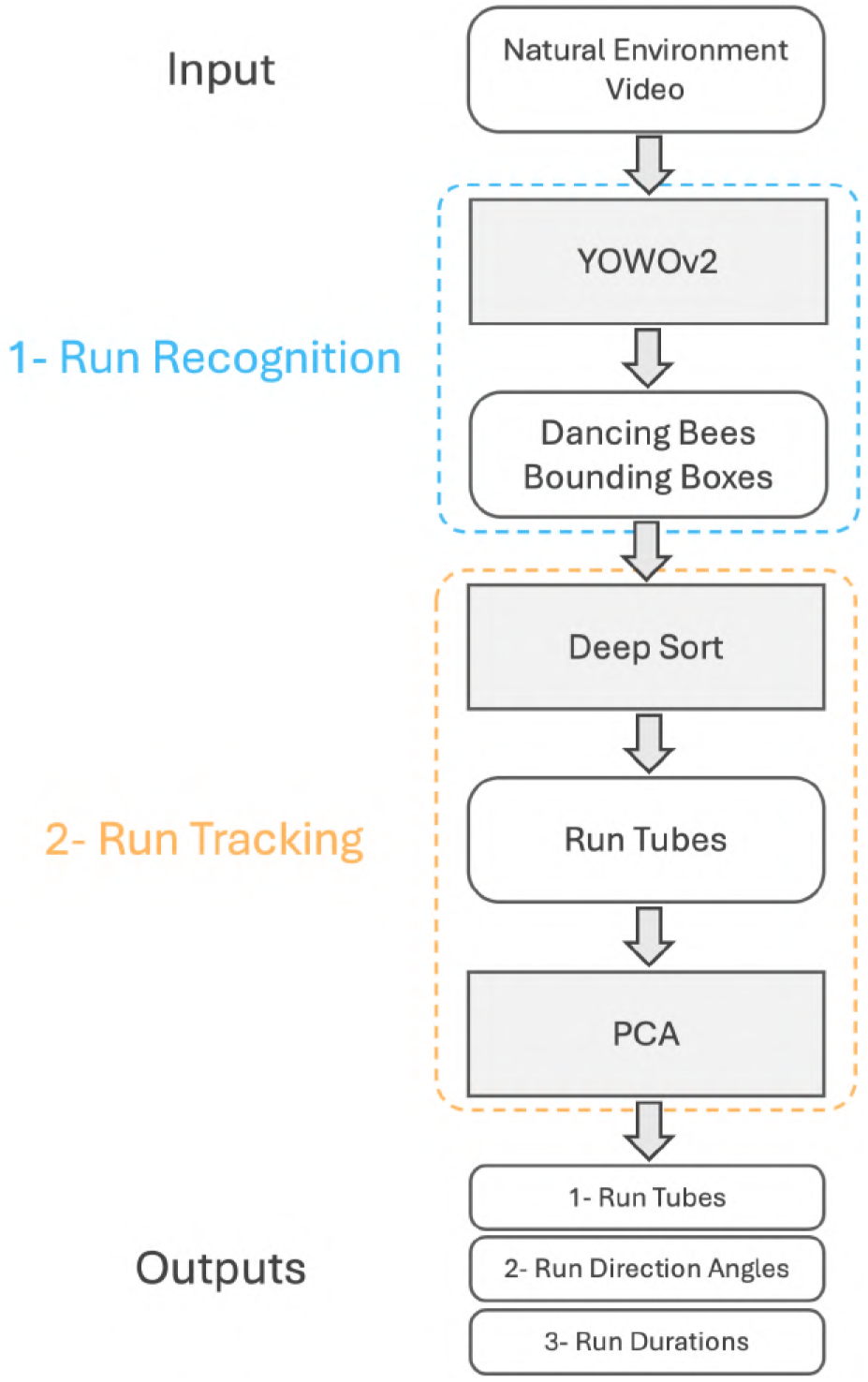
Architecture of our pipeline.

Recognition, tracking and parameters extraction are made for each waggle run individually.

#### 2.2.2 Run recognition

The run recognition module is based on the YOWOv2 video-based deep learning model. YOWOv2 is a detector that takes a video as an input and outputs detected bounding boxes for each of its frames (Yang & Dai, 2023). YOWOv2 is comprises of three parts: a backbone, a neck with a feature pyramid network and a detection head (Fig. 4). The backbone profits from combining both 2D and 3D backbones to take benefits of spatial and spatio-temporal features. Unlike previous approaches, using spatio-temporal features considerably increases the consistency of detections over time, reduces the number of false negatives and significantly reduces the number of missed detections. Furthermore, based on our research, a specific motions of a living organism like the waggle run is difficult to detect based on the information from one frame only. YOWOv2’s backbone was equipped with a modified version of the backbone of YOLOv7 as a 2D backbone and the efficient 3D CNN as a 3D backbone (Kopuklu et al., 2019; Wang et al., 2023). YOWOv2 can process different sizes of clips to produce detections. Given a video clip with *K* frames *V* = *{I*_1_*, I*_2_*, . . ., I_K_}*, where *I_K_* is the last frame of the current clip, *I_K_* is the input of YOWOv2’s 2D backbone and *V* is the input of its 3D backbone. 2D features are extracted at three different scales and 3D features are extracted and upscaled twice to match these three scales. In particular, the 2D backbone extracts two decoupled sets of features: classification features processed to focus on classifying objects within the frame and regression features processed to predict the position and size of the bounding boxes. Finally, the detection head merges 2D and 3D features through a channel encoder before outputting the final predictions using a classification branch and a box regression branch, without using anchor boxes. Outputs in our case are lists of bounding boxes of bees performing waggle runs for all frames of a video. In our research, only one class was considered, "waggle run bee", which could be extended in the future to detect return runs or other specific behaviors.

**Figure 4:**
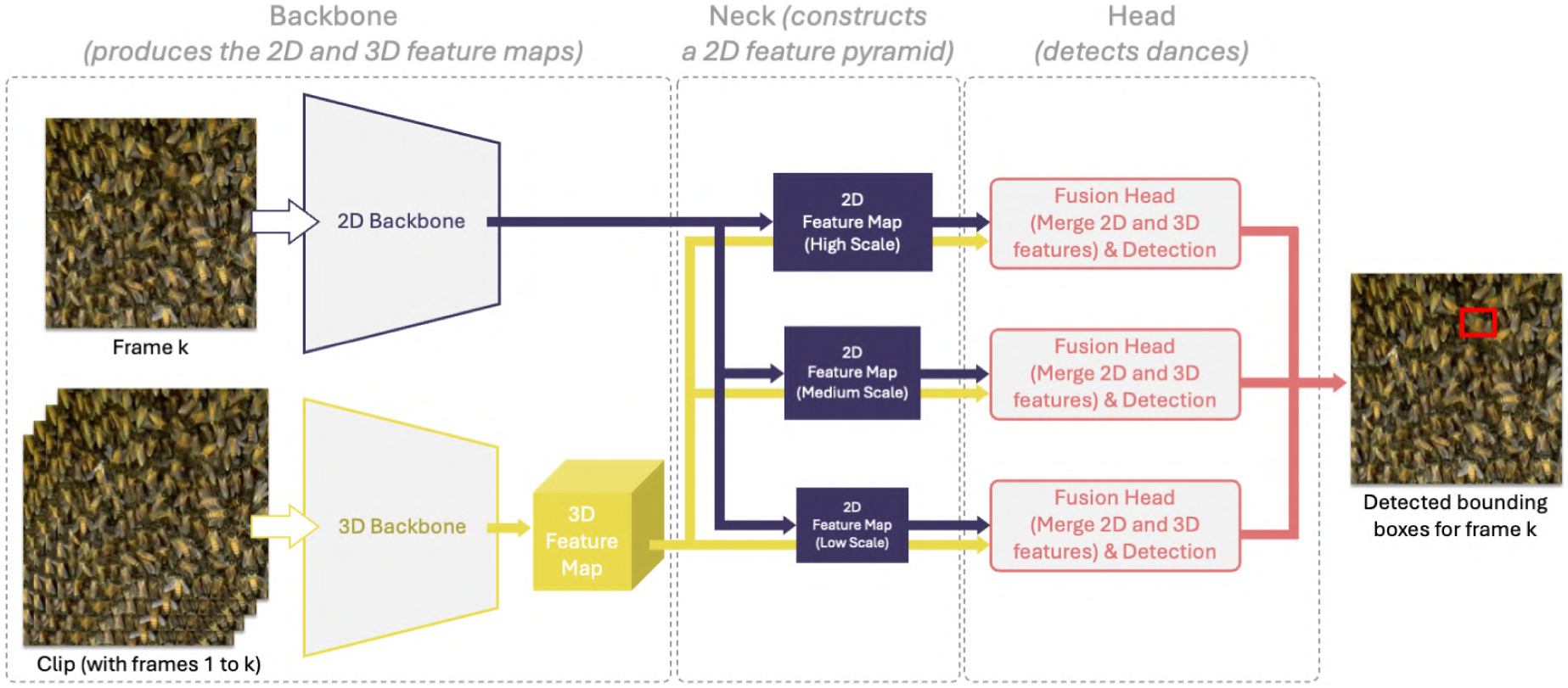
Simplified scheme of the architecture of YOWOv2.

The run recognition module was trained on the training dataset and its hyperparameters were tuned on the validation dataset using the built-in metrics frame AP and video AP and the downstream metrics from the run tracking module (Hou et al., 2017; Redmon et al., 2016). The metrics used for run recognition are defined as:

- Frame AP; the area under the precision-recall curve based on detections from frames. In our case, a detection at one frame is considered correct when the intersection-over-union(IoU) with the ground truth bounding box is greater than an IoU of 0.5.
- Video-AP; the area under the precision-recall curve of the action tubes predictions. In our case, a tube is considered correct when the mean of the IoUs over this tube with the ground truth tube is greater than an IoU of 0.3. A lower IoU was used for video AP to expand its range and facilitate the selection of the superior video AP value.

The run recognition module has two hyperparameters: the number of frames used as inputs for the 3D backbone (1, 2, 4, 8, 16 or 32) and the size and architecture of the YOWOv2 model, i.e. YOWOv2 nano which uses a shufflenet-v2 as a backbone in its 2D backbone or YOWOv2 tiny which uses the same backbone as YOLOv7 (Ma et al., 2018). We trained the model for 20 epochs for each set of hyperparameters and used the same loss function as in the original paper with AdamW as an optimizer and a cosine learning rate scheduler comprising one warm-up epoch (Loshchilov, 2017). Data augmentation was applied to inputs through random cropping, random flipping and random distortion.

#### 2.2.3 Run tracking

To decode waggle runs and map the indicated distance and direction of the food sources, two features are needed (Schürch et al., 2019):

- The linear trajectory of the bee’s movement pattern, which can be translated to the direction to the foraging site.
- The duration of the dance, which can be translated to the distance to the food source.

The run tracking module uses the deep learning tracking algorithm DeepSORT to temporally connect detected dances from frames and reconstruct waggle runs, in order to obtain tubes and calculate the desired features. To reduce false positives, only waggle runs that exceeded a threshold duration were selected for further analysis. The lowest boundary of this range was set to match the duration of the shortest waggle run of our training dataset. At this point, dance tubes were obtained and their durations were deduced from their number of frames. Then, a PCA was performed using the center point of detections within each waggle run as points (Maćkiewicz & Ratajczak, 1993). The first principal component of the PCA, corresponding to the largest eigenvalue, represents the direction of maximum variance in the data and was interpreted as the best-fit line for the waggle run’s trajectory. The angle of each waggle run relative to the vertical axis was determined from this main axis of the PCA. This angle was first computed using the standard PCA output from equation (eqn 1), and then transformed to match the coordinate system and orientation conventionally used in waggle run research (Fig. 5). These transformed angles became waggle run angles. Later, these waggle run angles can be transformed into real world "indicated directions" by adding sun’s azimuth during the dance. By using PCA in this way, we were able to efficiently determine the overall direction of each waggle run, even in the presence of small deviations or noise in the individual detection points. This method provides a robust estimate of the run’s orientation, which is crucial for accurate analysis in waggle run research contexts.

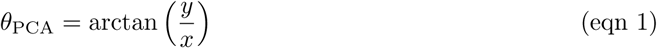

**Figure 5:**
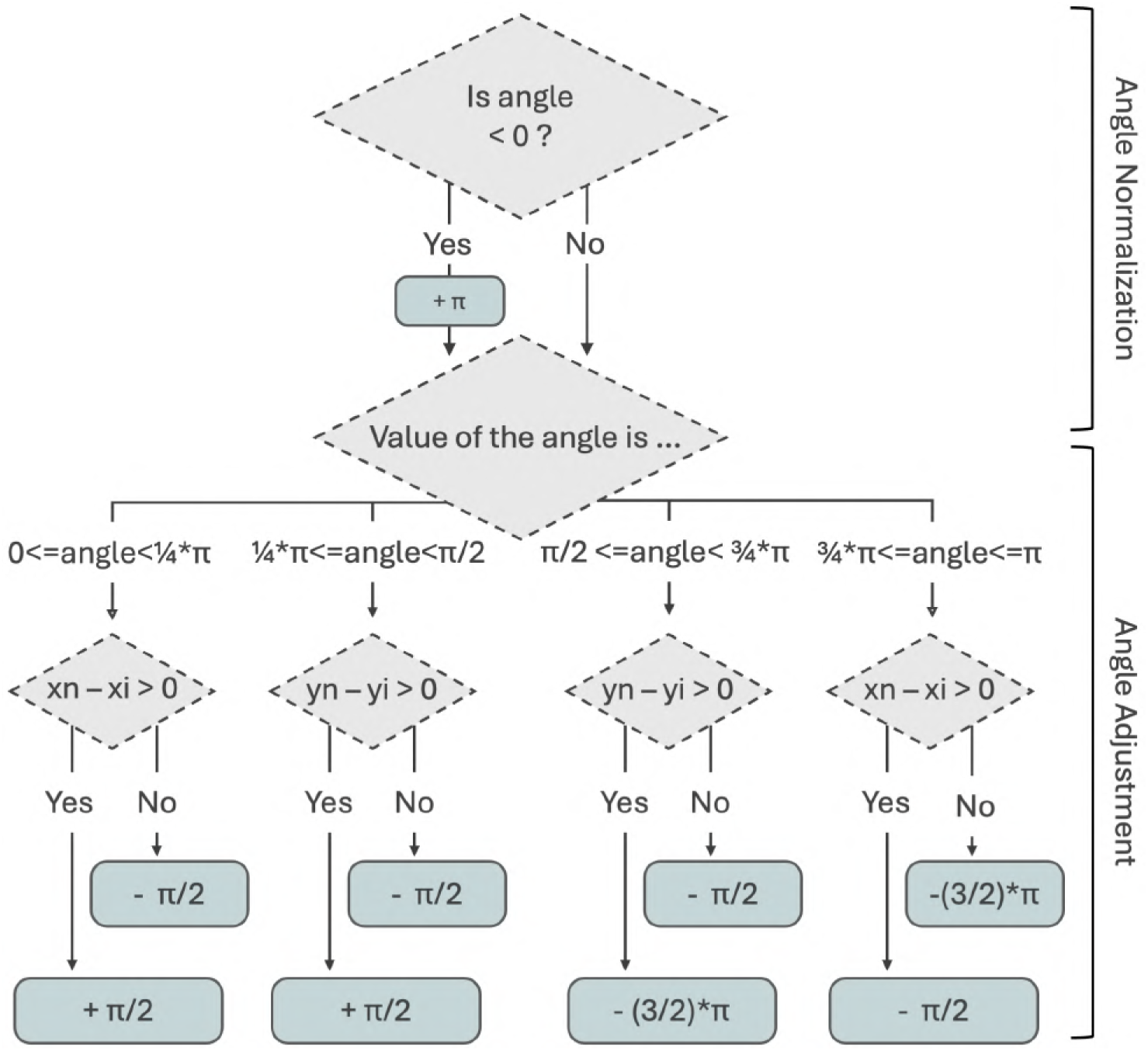
Correction pipeline for angles from PCA. (xi,yi) and (xn, yn) are respectively the coordinates of the first and last bounding boxes’ centers during the waggle run.

where *x* and *y* are the coordinates of the support vector of the first component of the PCA.

Four metrics were computed after the run tracking module: percentage of correctly detected waggle runs, angle RMSE (Root Mean Square Error) and duration RMSE (equation (eqn 2)), and duration bias (equation (eqn 3)), all compared to the actual values, represented by the manual annotation. Duration RMSE and angle RMSE measure the difference between ground truth and predicted values on correctly detected waggle runs. Duration bias gives insights about a potential systematic under-estimation or overestimation of a waggle run duration. The last metrics that we measured was the percentage of correctly detected waggle runs. It was measured without regard to the overlap between detection and ground truth tubes. Duration RMSE, angle RMSE and duration bias were computed without taking missed detection into account. Usually, a bee sequentially performs a series of very similar waggle runs for the same location as it recruits new foragers. For this reason, we considered that a few missed runs would not be a problem for the final analysis of the foraging sites: in most cases other waggle runs in the same series will point to the same area. Indeed, when analyzed, final angles and duration are typically averaged over all waggle runs from each individual. As a consequence, we estimated that the percentage of correctly detected waggle runs was the most important metric for hyperparameter tuning analysis.

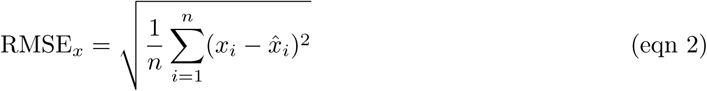

where:

- *x* represents the variable being measured (either *θ* for run angle or *τ* for run duration)
- *x_i_* is the ground truth value for the *i*-th measurement
- 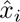 is the estimated value for the *i*-th measurement
- *n* is the total number of measurements

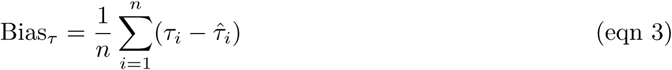

where *τ_i_* is the ground truth waggle run duration and 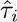 is the estimated waggle run duration.

### 2.3 Mapping of the runs

To visualize the practical application of our pipeline, a map of potential foraging sites is created based on the decoded waggle run targets. Using the colony’s known location as the central point, each waggle run target on the map is plotted. The sun’s azimuth was used to convert dance angles into real-world directions, while the model (i.e. equation of the linear slope of the dance distance calibration curve) from Kohl et al. (2020) helped to translate the waggle run durations into distances. Each decoded run is represented on the map as a colored marker, with different colors indicating different dancing individuals in the ground truth data. Then, as our model is not currently detected full dances but single runs individually, we used the DBSCAN clustering algorithm to group similar runs (based on angle and duration values), which likely represent multiple dances for the same food source (Ester et al., 1996). To provide an aggregated view, the mean position for each cluster is calculated and displayed. This mapping approach offers a visual representation of bee foraging patterns, allowing researchers to gain insights into the spatial distribution of food sources as communicated through waggle runs.

### 2.4 Generalizability of the model

In order too evaluate the model’s ability to accurately detect waggle runs across different contexts, and to gauge its potential for large-scale environmental studies in diverse ecosystems, we tested it on a novel dataset. This dataset consisted of 7 videos recorded from *Apis mellifera* beehives (Fig. 6) situated in Tokyo, Japan, captured over two different days in June 2018, totaling 35 minutes of footage. Importantly, the model had not been retrained on data from this specific month or on any *Apis mellifera* videos, ensuring that it had no prior exposure to this particular context.

**Figure 6:**
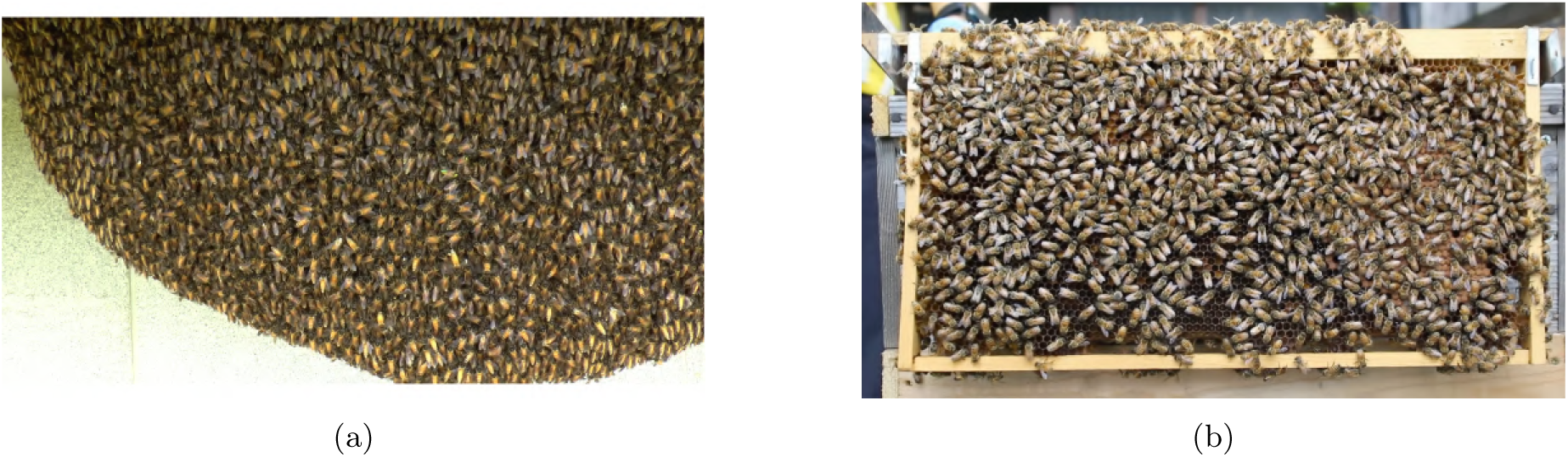
Comparison of the two types of hives we shot: a hive of *Apis dorsata* (a) and a hive of *Apis mellifera* (b).

### 2.5 Computational Resources and Performance Metrics

Training and inference of our pipeline were processed using a computer equipped with an Intel Core i9-10850K CPU, a NVIDIA GeForce RTX 3090 GPU and 64GB RAM. Table 1 shows the computational costs of the selected architecture over inference on the testing dataset, with a batch size of 1 and a video size of 448x448. Memory usage and GPU memory usage were computed using the python libraries psutil (https://pypi.org/project/psutil/) and GPUtil (https://github.com/anderskm/gputil). This is an indication for researchers who would like to run our code on their own hardware.

**Table 1:** Run recognition and run tracking computational costs on the testing dataset.

| Module | FPS | Memory Usage | GPU Memory Usage |
| --- | --- | --- | --- |
| Run Recognition Module | 16±0 | 2398±1 Mb | 1305±4 Mb |
| Run Tracking Module | 26±2 | 3265±1 Mb | 2228±23 Mb |

## 3 Results

In this work, we evaluated the capacity of proposed pipeline to accurately detect bees waggle runs and extract two critical parameters for waggle run analysis: duration and angle. We also ran experiments to define the best hyperparameters.

### 3.1 Hyperparameters tuning on the validation dataset

We first trained run recognition on the training dataset and computed frame AP and video AP for different sets of parameters on the validation dataset. Since temporal consistency is crucial to extract run duration, we selected accuracy values corresponding to the epoch with the best Video 0.3 AP obtained over the 20 training epochs. Frame AP values show how consistent are detected bounding boxes compared to the ground truth. From Table 2, we can see that Frame AP values globally increase with the K parameter, by 14% for YOWOv2 nano (K=1) and 6% for YOWOv2 tiny (K=32). The improvement is even more clear when looking at the video AP, a metric that shows the consistency of detected tubes. Video AP increased by 54% for YOWOv2 nano (K=1) and by 29% for YOWOv2 tiny (K=3). Video AP seems to significantly improve when changing architecture, from YOWOv2 nano to YOWOv2 tiny. It is the highest for YOWOv2 tiny and a K of 16, with a value of 0.85. The results indicate that higher values of K and larger network architectures enhance the prediction accuracy. However, although run recognition metrics give an idea of the consistency in the detection of dancing bees, they do not show how accurate are the features after their extraction.

**Table 2:** Run recognition and run tracking metrics on the validation dataset.

| Architecture | K | Frame<br>0.5 AP<br>↑ | Video<br>0.3 AP<br>↑ | Detected<br>Runs (%)<br>↑ | Duration<br>RMSE (frame)<br>↓ | Duration<br>Bias (radian) | Angle<br>RMSE (radian)<br>↓ |
| --- | --- | --- | --- | --- | --- | --- | --- |
| YOWOv2<br>nano | 1 | 0.83 | 0.48 | 79 | 30 | -20 | 0.14 |
|  | 2 | 0.89 | 0.56 | 86 | 26 | -15 | 0.14 |
|  | 4 | 0.92 | 0.70 | 90 | 22 | -14 | 0.12 |
|  | 8 | 0.94 | 0.70 | 86 | 18 | -11 | 0.12 |
|  | 16 | 0.94 | 0.73 | 93 | 22 | -14 | 0.13 |
|  | 32 | 0.95 | 0.74 | 90 | 24 | -15 | 0.13 |
| YOWOv2<br>tiny | 1 | 0.90 | 0.62 | 83 | 26 | -20 | 0.13 |
|  | 2 | 0.90 | 0.81 | 90 | 24 | -10 | 0.16 |
|  | 4 | 0.94 | 0.78 | 90 | 20 | -12 | 0.14 |
|  | 8 | 0.94 | 0.72 | 90 | 22 | -13 | 0.11 |
|  | 16 | 0.96 | 0.85 | 90 | 21 | -14 | 0.13 |
|  | 32 | 0.95 | 0.80 | 90 | 24 | -15 | 0.14 |

Metrics computed after run tracking on the validation dataset give insights into the accuracy in a practical application. Looking at Table 2, RMSE are lower K values greater than 1. For YOWOv2 nano, the greatest reduction of duration RMSE values (30 to 18) were produced by increased K from 1 to 8. For YOWOv2 tiny, duration RMSE decreases most when increasing K from 1 to 4. Angle RMSE values only vary slightly for different sets of parameters, ranging from 0.11 to 0.16 radian, representing an error of less than 10°. More generally, duration RMSE and angle RMSE do not give clear insight about which hyperparameters to choose. As run parameters are averaged over waggle runs before decoding, the error in decoding waggle dances to real world coordinates will decrease with the increasing number of correctly detected consecutive waggle runs. Given that, and the difficulty to draw conclusions from duration and angle RMSE on our dataset, we considered that the percentage of detected waggle runs was the most important metric to analyze. This value significantly increases from K of 1 to greater values for K, for both architectures. There was an obvious added value in using more than one frame which highlighted the importance of using video-based features for the detection of specific living organisms behaviors. However it does not significantly improve between YOWOv2 nano and tiny. One possible reason could be that the training dataset is not large enough to help a bigger architecture to extract generalized patterns in the data. The biggest percentage of detected waggle runs is achieved by YOWOv2 nano with a K of 16.

Results from both the run recognition and the run tracking modules motivated the choice of YOWOv2 nano with a K of 16 as hyperparameters. Duration bias naturally also has an improvement with greater K values. We observe that the bias is always negative, regardless of the set of parameters used. This result highlights the consistent overestimation of the network. In practice, every waggle run tube is detected with extra bounding boxes before and after the waggle run. Since duration bias achieved a value of -14 for this configuration, we applied a correction factor of the same value on output durations. The correction factor was determined on the validation dataset and assessed on the testing dataset.

### 3.2 Final assessment on the testing dataset

Results were consistent with the ones obtained on the validation dataset. Our pipeline achieved values of 0.97 for frame 0.5 AP and 0.68 for video 0.3 AP. Video 0.3 AP is lower than the value for the validation dataset but it seems to have a low impact on run tracking metrics. Duration RMSE achieved a value of 23 frames, a similar to the validation dataset scores. Angle RMSE followed the same direction, achieving 0.14 radian, when the correction factor was introduced. Since we used 50-fps videos, this represents an error of almost half a second. In addition, comparisons of ground truth and predicted angles and durations on the testing dataset are shown in Fig. 7. Angle points align almost perfectly with the identity line, achieving a R-square of 0.98, highlighting a high precision of our predictions. Duration points however show more variability and random errors around the identity line, with a R-square of 0.18. This is due to the difficulty of YOWOv2 to detect the real start and end of runs. The duration plot shows two points particularly far from the identity line and removing them would increase the R-square to 0.53. Still, the scatter plot is centered around the line and no longer shows systematic overestimation, with the incorporation of the correction factor. The aggregation of some waggle runs into cluster-like groups along the identity line is explained by the fact that the dataset is built around waggle runs that come from the same waggle dances. Finally, a example of a detection sequence and some comparisons of ground truth and predicted trajectories from the testing dataset are shown in Fig. 8 and Fig. 9 and demonstrate the proximity of ground truth and predicted trajectories. All dataset trajectories are shown in Appendix A.

**Figure 7:**
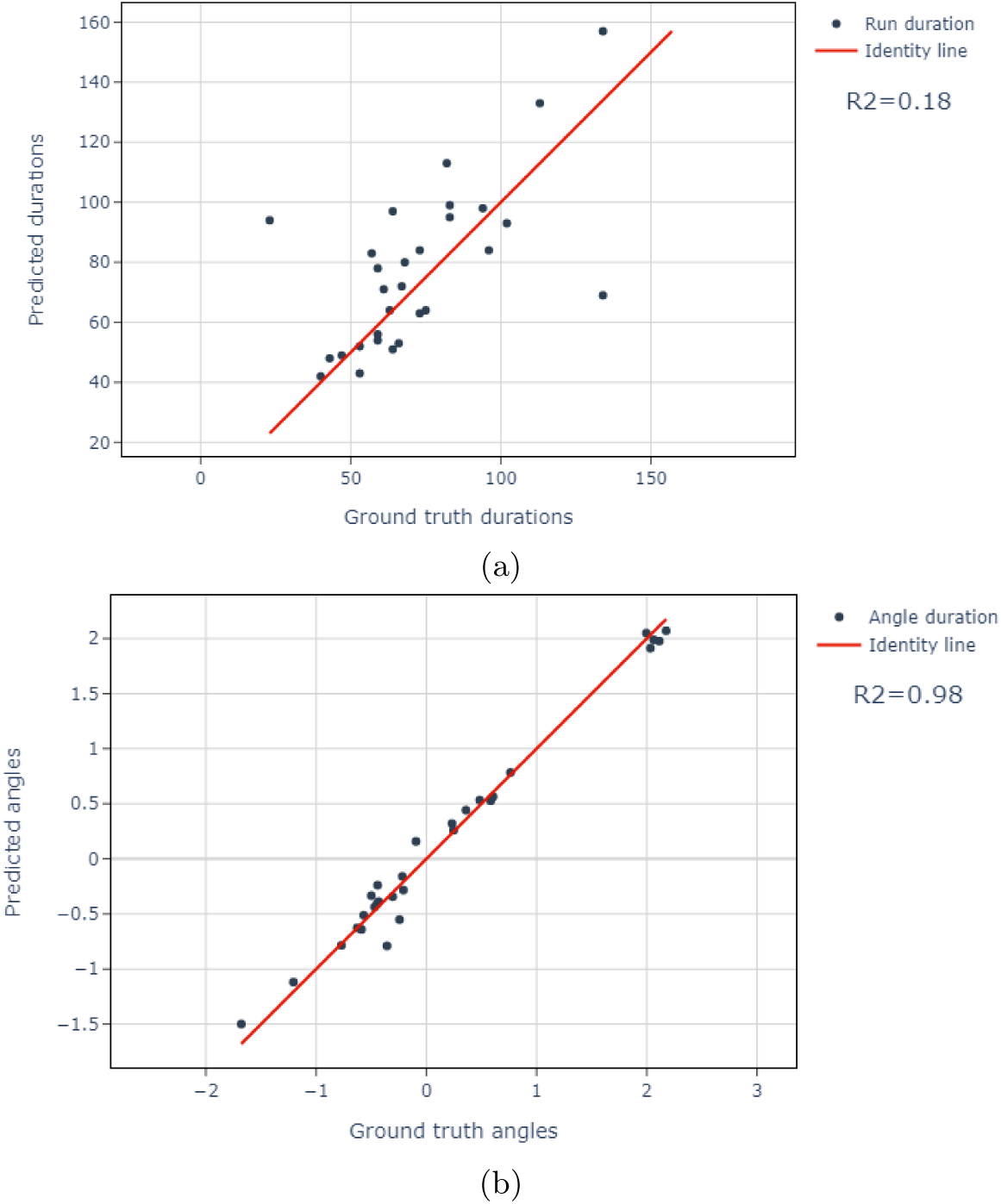
Comparisons of ground truth and predicted run features for YOWOv2 nano with K=16 and a correction factor of -14 on the testing dataset. (a) duration, (b) angle.

**Figure 8:**
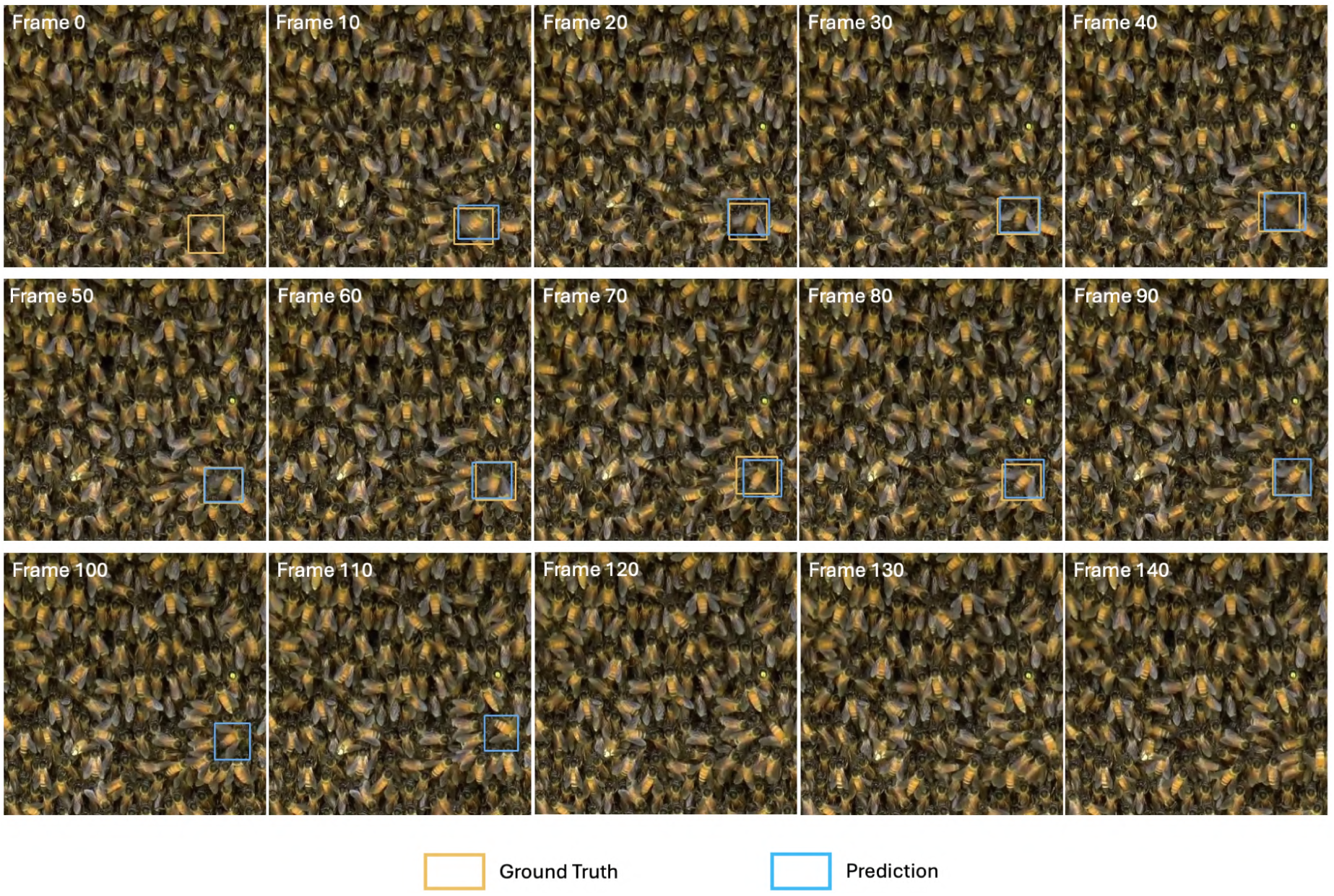
Comparison of YOWOv2 predictions to the ground truth for one video.

**Figure 9:**
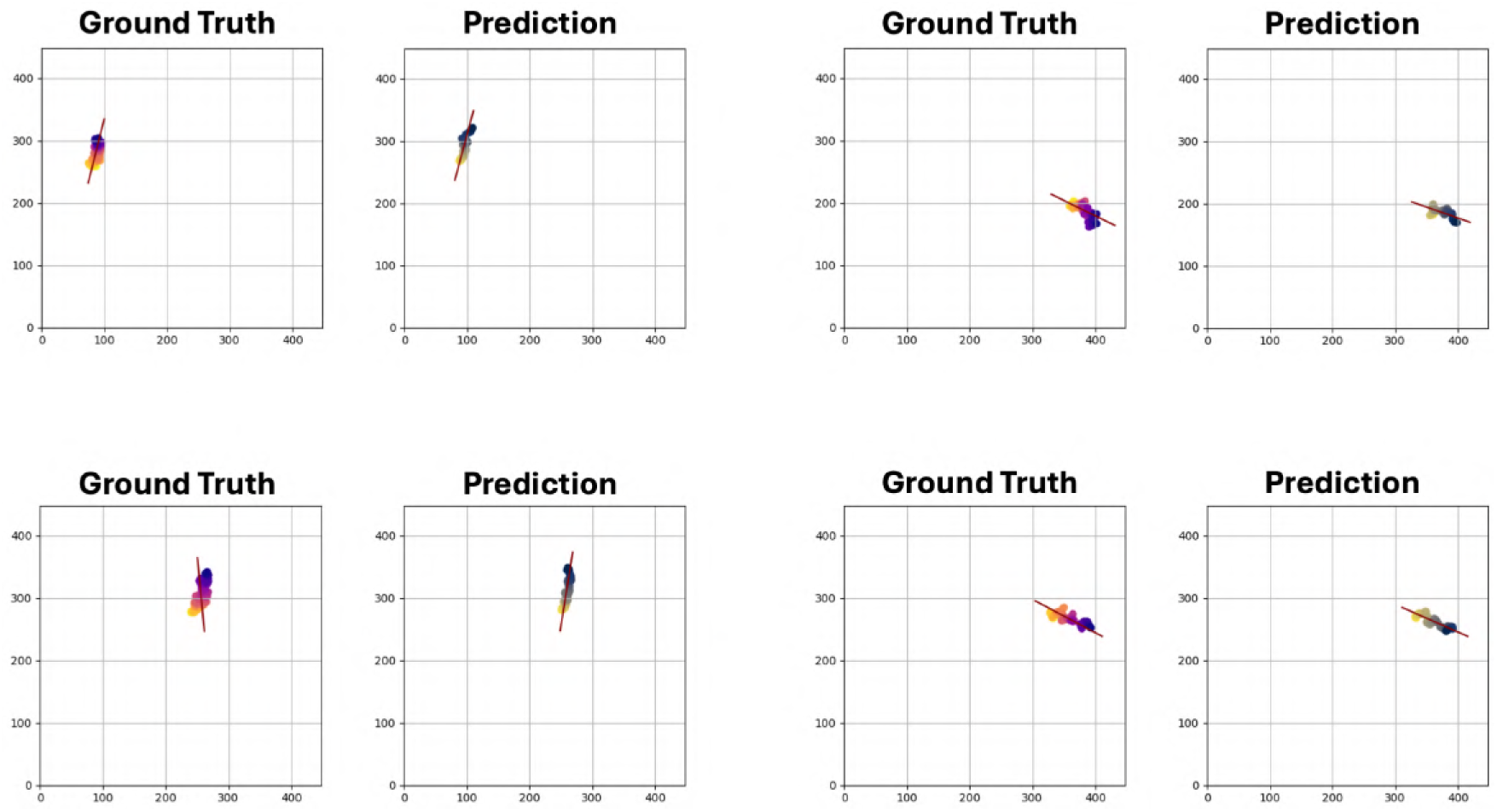
Comparison of ground truth and predicted trajectories for YOWOv2 nano with K=16 on the testing dataset. In each plot, points are bounding box centers for different frames and the line shows the orientation of the angle of the run obtained from PCA. Time on the graph progresses in the same direction as the transition from light yellow to dark blue. x-axis and y-axis are the coordinates in terms of pixels.

### 3.3 Mapping of potential foraging sites

Fig. 10 illustrates the mapping of analyzed waggle runs from the testing dataset, showing the potential of proposed pipeline for understanding bee foraging behavior in a spatial context.

**Figure 10:**
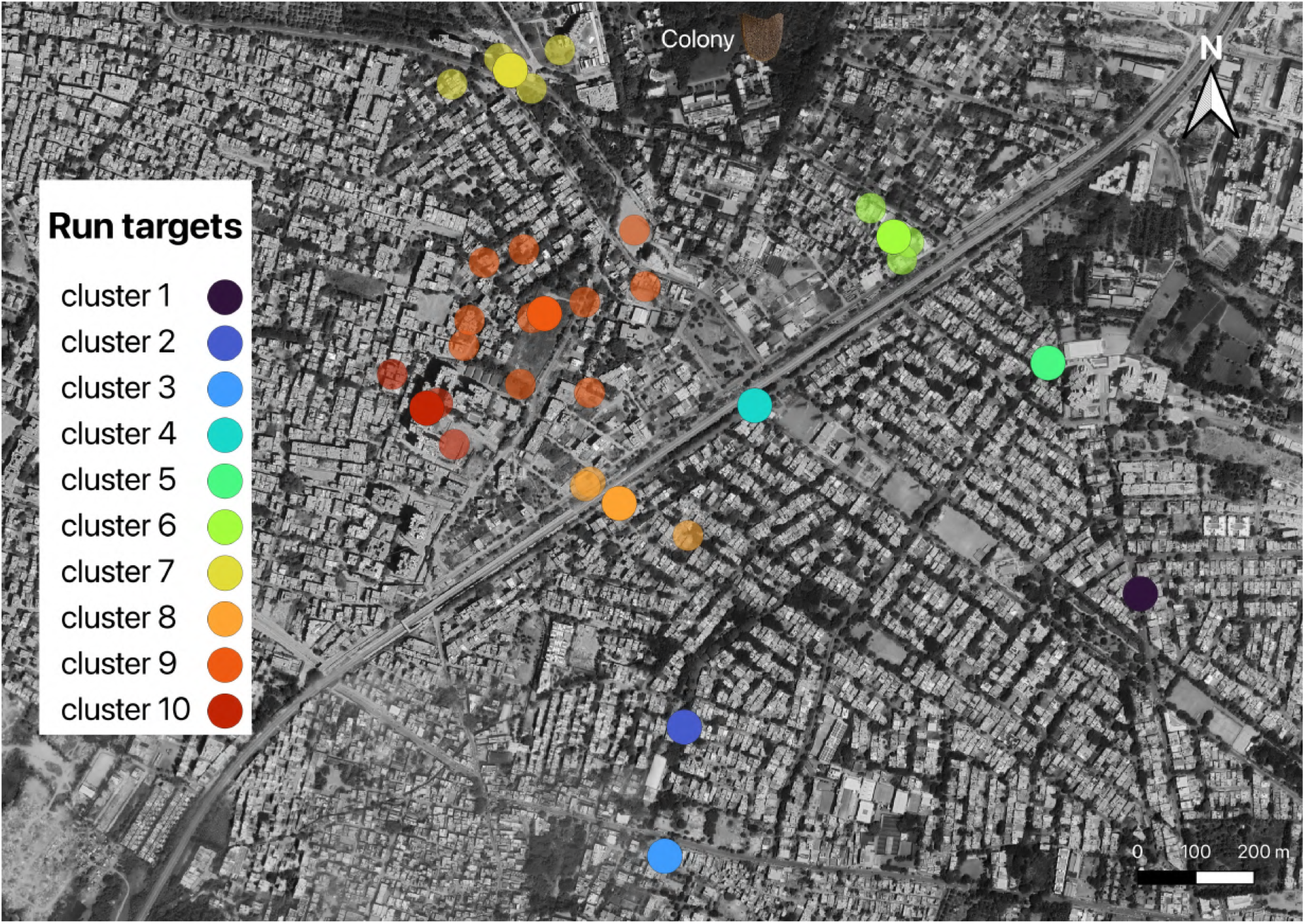
Map of the translated targets of the runs predicted by our model on the testing dataset. Transparent dots are the decoded positions and opaque dots are the means, averaged within a cluster.

### 3.4 Model performance across new contexts

We evaluated the model’s performance on a new dataset consisting of videos recorded from *Apis mellifera* hives. In total, we identified 140 waggle dances in the dataset, of which 70 were successfully detected by the model, achieving 50% of detected dances. The detection rate was lower than the one achieved on the testing dataset featuring videos from *Apis dorsata* hives. However, this shows that the model has a transferability potential to new recording conditions since both the environment (including factors such as hive type, lighting, etc.) and the bee species (meaning different body characteristics, such as a smaller size or a different body pattern) differed.

## 4 Discussion

Previous approaches to analyze bee waggle runs largely relied on manual observation or simpler image processing techniques, which present significant limitations in terms of scalability, efficiency, and applicability to analyse video recordings from hives under natural conditions. Manual decoding of waggle runs is time-consuming, labor-intensive, and subject to human error and variation among different annotators. Earlier automated methods, such as those developed by Rau (2014) and Wario et al. (2015, 2017), while innovative, often required specialized observation hives, controlled lighting conditions, and high-speed cameras, limiting their use in natural settings. These methods also frequently relied on predefined thresholds and frequency-based detections, which may not generalize well across different environments or bee species. In contrast, our deep learning-based pipeline offers several key advantages. It can operate effectively on videos captured in natural conditions using standard Full HD cameras, without the need for controlled lighting or specialized hives. The use of YOWOv2, a video-based deep learning model, allows for more robust detection by leveraging spatio-temporal features, significantly improving the consistency and accuracy of waggle run detection. Our method achieves comparable or better detection rates (up to 93% of runs detected) while working under more challenging, real-world conditions. The method’s capacity to operate effectively with standard Full HD cameras and without controlled lighting conditions represents a major advancement as it does not require a context-specific parameter tuning.

Certain limitations and challenges were encountered during the study. Firstly, the relatively small size of our dataset (127 training videos) may limit the model’s ability to adapt across a wider range of recording conditions for the moment. In comparison, the original YOWOv2 model was trained on the AVA dataset, consisting of 430 15-minute videos of various human activity scenes, and on the UCF101-24 dataset consisting of a subset of UCF101 including 3207 videos of human action videos collected from Youtube (Gu et al., 2018; Soomro et al., 2012). However, we did apply our model on videos featuring very different characteristics and still achieved a 50% run detection rate with *Apis mellifera* hives, without any specific training on this specie and this context. This in itself is a promising result toward a holistic model that could handle various recording parameters if trained with variations in datasets. The small size of the datasetwas particularly evident in the difficulty of precisely determining the start and end points of waggle runs, leading to some inaccuracies in duration measurements (duration RMSE of 23 frames). The use of 50 fps videos, while sufficient for detection, may also have contributed to these timing discrepancies. Additionally, the study focused primarily on *Apis dorsata*, potentially limiting the model’s immediate applicability to other bee species. These limitations affected our results by introducing some errors in duration measurements (R-square of 0.18). To address these challenges, we implemented a correction factor for duration and emphasized the importance of averaging measurements over multiple runs. Due to the size of our validation and testing datasets, it is difficult to run a deeper error analysis on duration. On the other hand, angle predictions were highly accurate (R-square of 0.98) using only a basic method: a PCA on bounding box centers. There is also room for improvement by using a specialized bee’s body part machine learning detector. Previous papers introduced such insect pose estimation methods but they were mostly conducted under controlled laboratory environment (Duan et al., 2017; Pereira et al., 2019; Wen et al., 2015). Additionally, deploying such a method would require heavier computation and greater data annotation efforts to achieve a minor improvement in angle prediction accuracy. Our method clearly demonstrates that deep learning has the potential of studying insects in its natural environments while alleviating the human labor problem with the least effort to date.

Secondly, in our results, inconsistency between values produced by our pipeline and the ground truth may also arise in part from human error in annotating the ground truth waggle run duration and angle. Given the same video, different annotators may give different annotations. Using a higher frame-rate than the 50fps used in our study would help the annotator to identify the beginning and the end of each waggle run more precisely. From the pipeline side, the error might also be reduced with a higher frame-rate. However, increasing the frame-rate would also increase the annotation time. Future studies may compare AI predictions to human annotations from various annotators, with various frame-rates, in feeder experiments. Regarding this comparison, we want to bring two elements to the attention of the reader. First, individual bees usually perform multiple waggle runs pointing at the same direction. The number of repeated waggle runs can be translated to desirability of the food source (Grüter & Farina, 2009; Seeley et al., 2000). Thus, averaging both predicted angles and predicted durations over dances has the potential to improve the spatial accuracy of the translations. The number of detected waggle runs (up to 93%) by our model is promising and must be kept high since it could improve the accuracy of the predictions. The second element is that there could be an intra-bee variation in the waggle runs performed by the same bee. The question of the variation being an adaptation or the result of a limit from the bee is an open discussion (Preece & Beekman, 2014). Also, a bee might be obstructed by other bees while dancing, leading to an imperfect movement. Thus, it is not possible to directly translate human or machine prediction errors to spatial errors without having the real target information in the training dataset.

The accurate detection and analysis of bee waggle runs, as enabled by our deep learning pipeline, offers profound implications for understanding bee communication, behavior, and ecology. This automated approach allows for continuous, large-scale monitoring of colony foraging patterns, providing unprecedented insights into how colonies interact with their environment. Such data is crucial for mapping critical foraging sites, understanding temporal dynamics of resource utilization, and identifying the impacts of environmental changes on bee behavior. In urban environments like Bengaluru, where this study was conducted, this information can guide conservation efforts by highlighting areas of importance for bee populations and informing bee-friendly urban planning initiatives. For agriculture, accurate waggle run analysis can reveal preferred crop species and optimal planting locations to enhance pollination services. It can also help in assessing the effectiveness of pollinator-friendly farming practices. Moreover, by enabling the study of bee communication in natural conditions, our method can contribute to a deeper understanding of how factors like urbanization, climate change, and habitat fragmentation affect bee colonies’ foraging strategies and overall health. This knowledge is vital for developing targeted conservation strategies and for maintaining the crucial ecosystem services provided by bees. The scalability of our approach also opens up possibilities for comparative studies across different bee species and geographical locations, potentially revealing broader patterns in bee behavior and adaptation to varied environments.

Building on the success of our current study, several exciting avenues for future research emerge. Expanding the scope of behavioral analysis beyond waggle runs to include other important bee behaviors, such as trophallaxis, grooming, shaking or guarding, as well as bee health studies in different environment such as through toxicovigilance, could provide a more comprehensive understanding of colony dynamics (Olivares-Pinto et al., 2024). Adapting the pipeline to work with different bee species, particularly in comparing communication methods across *Apis* species, could yield valuable evolutionary insights. To enhance the model’s performance and generalizability, future data collection efforts should focus on capturing videos across a wider range of environmental conditions, seasons, and geographical locations. This expanded dataset would not only improve the accuracy of duration measurements but also increase the model’s robustness to varying lighting conditions and backgrounds. Integration of multi-modal data could significantly enrich the analysis; for instance, combining video data with acoustic sensors to capture waggle run vibrations, or with environmental sensors to correlate waggle run patterns with temperature, humidity, or air quality. Such multi-modal approaches could provide a more holistic view of the factors influencing bee behavior. Another promising direction is the development of real-time processing capabilities, enabling immediate analysis and response in field conditions. This could be particularly valuable for ecological monitoring and conservation efforts. Furthermore, advancing the model to identify individual bees consistently across multiple waggle runs could allow for tracking of forager recruitment and the spread of information within colonies. As deep learning techniques continue to evolve, incorporating more advanced architectures or unsupervised learning methods might lead to the discovery of previously unrecognized patterns in bee behavior. Ultimately, these advancements have the potential to revolutionize our understanding of social insect communication, contribute to more effective conservation strategies, and inform sustainable agricultural practices in the face of global environmental challenges.

## 5 Conclusion

This research has successfully demonstrated the potential of using a deep learning-based pipeline to analyze the waggle runs of *Apis dorsata* and its generalizability to another bee specie in natural conditions which has never been successfully attempted prior to this. Our innovative approach not only automates the detection of bee performing the waggle dance but is also able to decode the dance and thus able to precisely measure the direction and duration, which are crucial to understand the direction and location of the bee forage and their communication methods. The ability to process video data autonomously allows for the study of bee foraging behavior at a scale that was unimaginable with traditional manual methods. Furthermore, the implications of this study extend beyond the scientific understanding of bee communication. By elucidating the foraging patterns of bees and the impacts of urbanization on their behavior, this research could contribute to conservation efforts, aiding in the design of bee-friendly urban planning initiatives. This is critical in preserving biodiversity and maintaining the ecosystem services that bees provide, particularly in rapidly urbanizing regions. For future research, enhancing the pipeline by integrating more diverse datasets, including different species and environmental conditions, could further improve the robustness and applicability of the model while opening the possibility to study different behaviors. Additionally, exploring real-time processing capabilities could enable the immediate application of this research in field conditions, providing beekeepers and conservationists with a powerful tool for monitoring and supporting bee populations.

## Acknowledgements

We thank Kozue Wada from the Laboratory of Field Phenomics, University of Tokyo, for her crucial help with image annotation and Shuai Xiang from the Laboratory of Field Phenomics, University of Tokyo, for his advice concerning the coding experiment. Smitha Krishnan would like to thank all funders who supported her through Nature-Positive Solutions, Transforming Agrifood Systems in South Asia (TAFSSA) and Agroecology Initiative. She acknowledges and appreciates the One CGIAR donors. This work was part of the initiative "Exploring and managing human–bee conflict in Asian cities using AI" supported by Google in the framework workshop "AI for social good."(https://www.cgiar.org/funders/). Sylvain Grison was supported by JST SPRING, Grant Number JPMJSP2108. in 2022. Rajath Siddaganga was supported by a grant from the Science and Engineering Research Board (SERB) CRG/2019/005984. Axel Brockmann was supported by NCBS-TIFR institutional funds (No. 12P4167) and the Department of Atomic Energy, Government of India (No. 12-R&D-TFR-5.04–0800 and 12-R&D-TFR-5.04–0900).

## Author Contributions

Wei Guo, Axel Brockmann, Smitha Krishnan led the study; Sylvain Grison, Wei Guo, Pieter M. Blok, James Burridge designed the deep learning pipeline; Rajath Siddaganga, Shrihari Hedge, Sylvain Grison collected the data; Rajath Siddaganga, Sylvain Grison analyzed the data, Sylvain Grison, Wei Guo, Axel Brockmann led the writing of the manuscript. All authors contributed critically to the drafts and gave final approval for publication.

## Data Availability

Our code is open-source and available on GitHub at https://github.com/UTokyo-FieldPhenomics-Lab/DeepWDT/tree/main. By sharing our work, we aim to facilitate the efficient and straightforward analysis of more waggle runs by the research community.

## Appendix A: Comparison of ground truth and predicted trajectories on the testing dataset

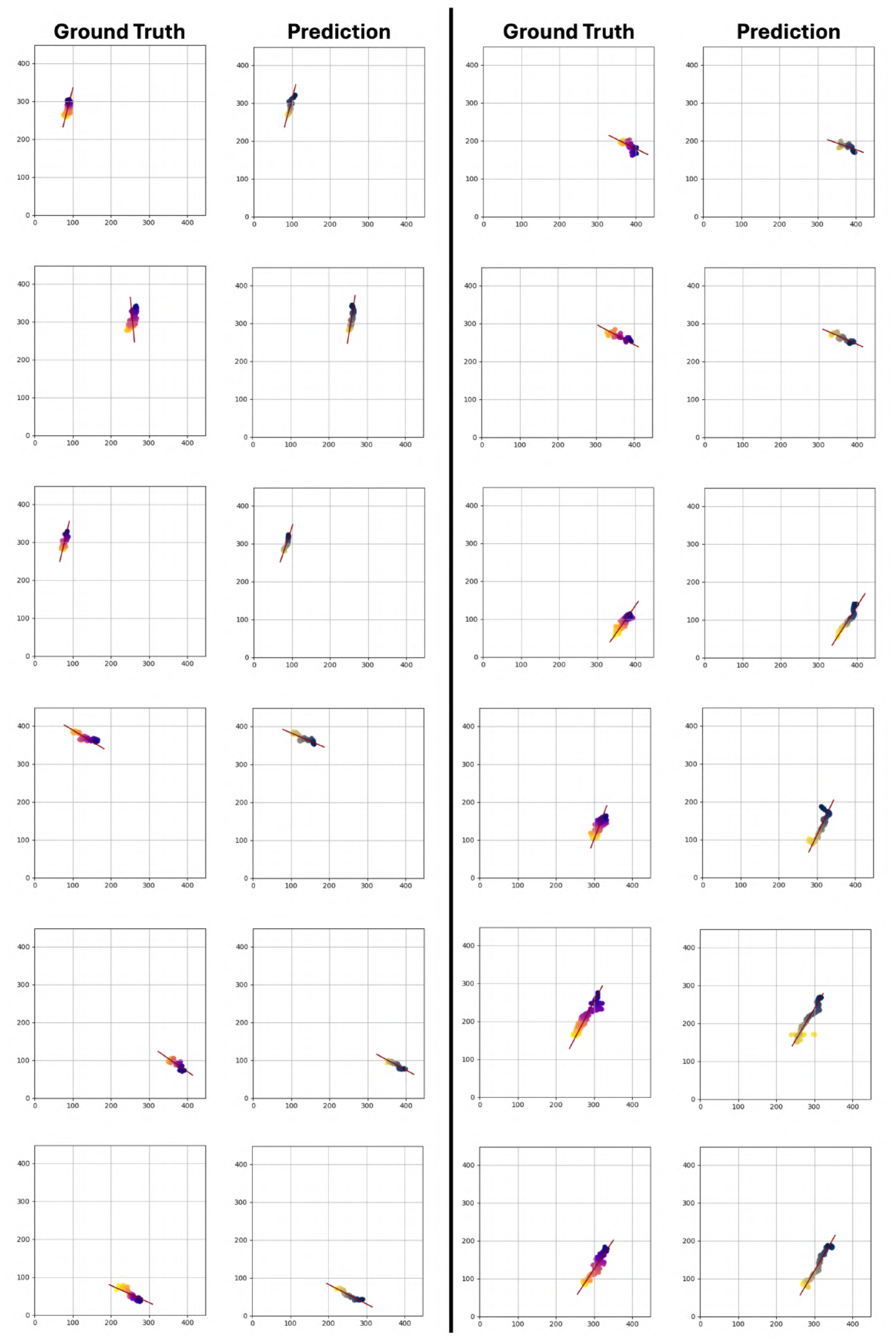
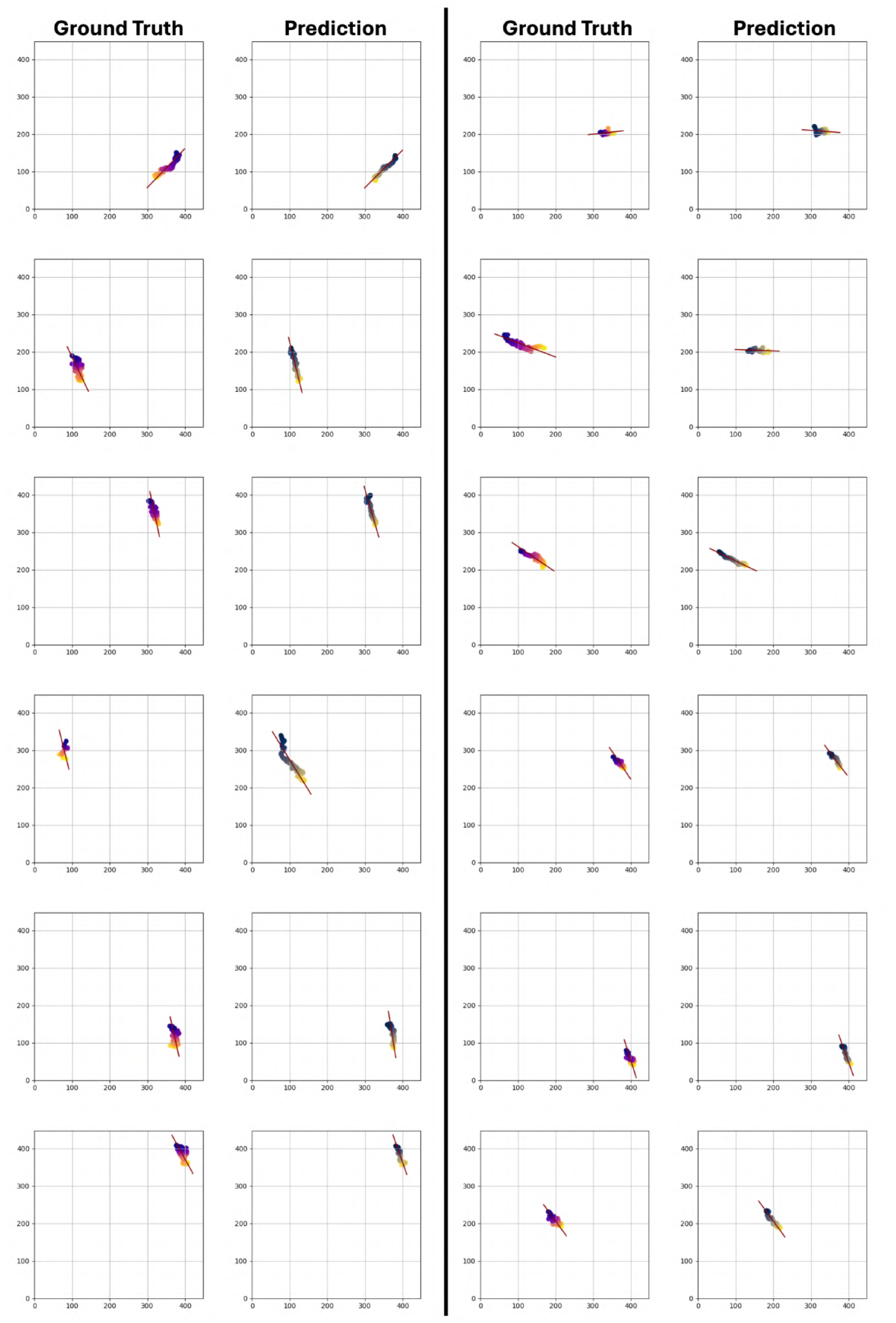
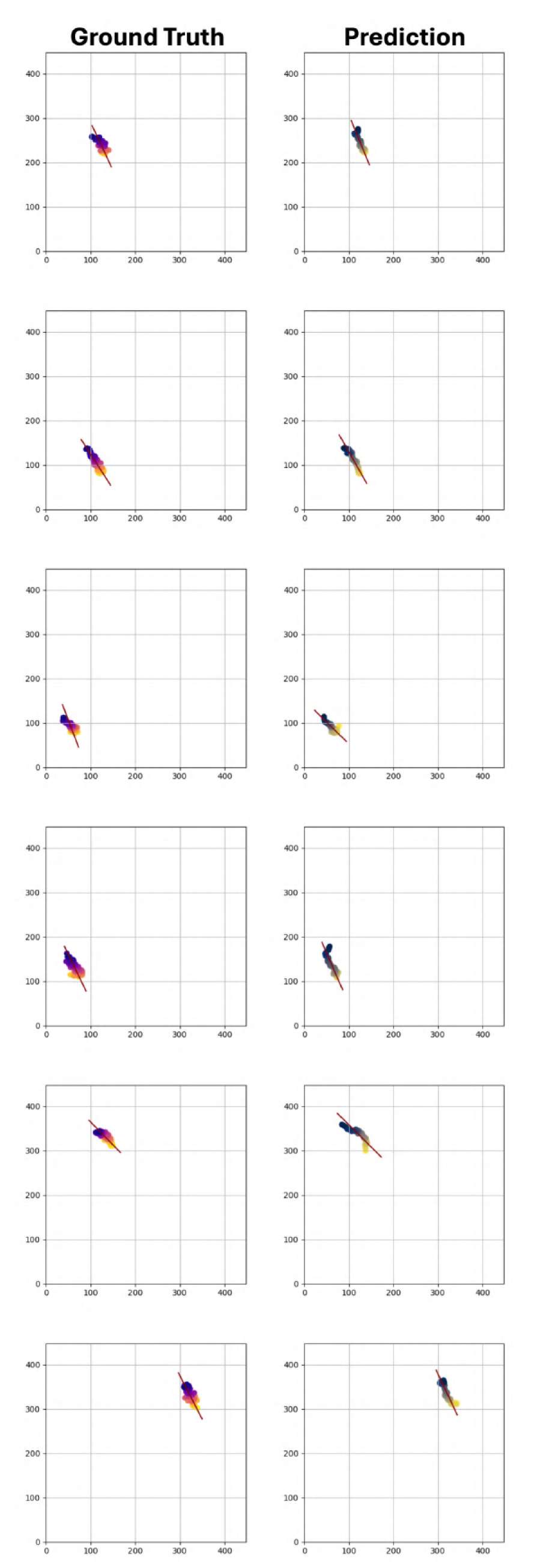

